# Chromosome-scale comparative sequence analysis unravels molecular mechanisms of genome evolution between two wheat cultivars

**DOI:** 10.1101/260406

**Authors:** Anupriya Kaur Thind, Thomas Wicker, Thomas Müller, Patrick M. Ackermann, Burkhard Steuernagel, Brande B.H. Wulff, Manuel Spannagl, Sven O. Twardziok, Marius Felder, Thomas Lux, Klaus F.X. Mayer, International Wheat Genome Sequencing Consortium, Beat Keller, Simon G. Krattinger

## Abstract

**Background:** Recent improvements in DNA sequencing and genome scaffolding have paved the way to generate high-quality *de novo* assemblies of pseudomolecules representing complete chromosomes of wheat and its wild relatives. These assemblies form the basis to compare the evolutionary dynamics of wheat genomes on a megabase-scale.

**Results:** Here, we provide a comparative sequence analysis of the ~700-megabase chromosome 2D between two bread wheat genotypes – the old landrace Chinese Spring and the elite Swiss spring wheat line ‘CH Campala *Lr22a*’. There was a high degree of sequence conservation between the two chromosomes. Analysis of large structural variations revealed four large insertions/deletions (InDels) of >100 kb. Based on the molecular signatures at the breakpoints, unequal crossing over and double-strand break repair were identified as the evolutionary mechanisms that caused these InDels. Three of the large InDels affected copy number of NLRs, a gene family involved in plant immunity. Analysis of single nucleotide polymorphism (SNP) density revealed three haploblocks of ~8 Mb, ~9 Mb and ~48 Mb with a 35-fold increased SNP density compared to the rest of the chromosome.

**Conclusions:** This comparative analysis of two high-quality chromosome assemblies enabled a comprehensive assessment of large structural variations. The insight obtained from this analysis will form the basis of future wheat pan-genome studies.

## Background

Bread wheat (*Triticum aestivum*) was the most widely grown cereal crop in 2016. It serves as a staple food for over 30% of the world’s population and provides ~20% of the globally consumed calories [1]. Wheat is a young allopolyploid species with a genome size of 15.4-15.8 Gb, of which more than 85% is made up of highly repetitive sequences [2]. The allopolyploid genome arose through two recent, natural polyploidization events that involved three diploid grass species. The first hybridization event occurred 0.58 to 0.82 million years ago [3] between the A-genome donor wild einkorn (*T. urartu*) and a yet unidentified B-genome donor that was a close relative of *Aegilops speltoides.* This hybridization created wild tetraploid emmer wheat (*Triticum turgidum* ssp. *dicoccoides;* AABB genome) [4]. A second natural hybridization between domesticated emmer and wild goatgrass (Ae. *tauschii;* DD genome) resulted in the formation of hexaploid bread wheat (AABBDD genome) around 10,000 years ago [5]. The domestication of tetraploid emmer and the limited number of hybridization events with *Ae. tauschii* represent bottlenecks that resulted in a significant reduction of genetic diversity within the bread wheat gene pool. Natural gene flow between bread wheat and its wild and domesticated relatives as well as artificial hybridizations with diverse grass species partially compensated for this loss in diversity [3, 6].

The size, repeat content and polyploidy of the bread wheat genome have represented major challenges for the generation of a high-quality reference assembly. The first ‘early’ whole genome assemblies of hexaploid wheat and its diploid wild relatives were based on short-read sequencing approaches. These assemblies provided an insight into the gene space of wheat, but they were highly-fragmented and incomplete [7-10]. The first notable high-quality sequence assembly of wheat was produced from the 1-gigabase chromosome 3B of the hexaploid wheat landrace Chinese Spring. For this, 8,452 ordered bacterial artificial chromosomes (BACs) were sequenced and assembled, which resulted in a highly contiguous assembly (N50 = 892 kb) [11, 12]. More recent whole-genome shotgun assemblies had improved contiguousness compared to the ‘early’ assemblies (N50 = 25 – 232 kb) [13-15], but they still did not allow to compare the structure of wheat chromosomes on a megabase-scale.

Several recent technological and computational improvements however provided a basis to generate *de novo* assemblies of complex plant genomes with massively improved scaffold lengths and completeness. These advancements included (i) the integration of whole-genome shotgun libraries of various insert-sizes [16] or the use of long-read sequencing technologies such as single-molecule real-time sequencing (SMRT) [17] or nanopore sequencing [18], (ii) the improvement of scaffolding by using chromosome conformation capture technologies [19-23] or optical maps [24] and (iii) the improvement of assembly algorithms [4]. Chinese Spring is an old landrace that was selected for sequencing because it was used in a number of cytogenetic studies, which has resulted in the generation of many important genetic resources from this wheat line, including chromosome deletion lines [26] and aneuploid lines [27].

The understanding of the genetic variation will provide an insight into wheat genome evolution and its impact on agronomically important traits. The continuum of genetic variation ranges from single nucleotide polymorphism (SNPs) to megabase-sized rearrangements that can affect the structure of entire chromosomes [28]. Due to the absence of high-quality wheat genome assemblies, previous comparative analyses were limited in the size of structural rearrangements that could be assessed and typically, structural variants of a few base pairs up to several kb were analysed [29, 30]. Consequently, a comprehensive assessment of the extent of large structural rearrangements and their underlying molecular mechanisms is still lacking.

Here, we report on a chromosome-wide comparative analysis of the ~700 Mb chromosome 2D between the two hexaploid wheat lines Chinese Spring and ‘CH Campala *Lr22a*’. ‘CH Campala *Lr22a*’ is a backcross line that was generated to introgress *Lr22a*, a gene that provides resistance against the fungal leaf rust disease, into the genetic background of the elite Swiss spring wheat cultivar ‘CH Campala’ [31]. We previously generated a high-quality *de novo* assembly from isolated chromosome 2D of ‘CH Campala *Lr22a*’ by using short-read sequencing in combination with Chicago long-range scaffolding [32]. The resulting assembly had a scaffold N50 of 9.76 Mb. In particular, the focus of our study was on the identification and quantification of large structural variations (SVs). The comparative analysis of the 2D chromosome showed a high degree of collinearity along most of the chromosome, but also revealed SVs such as InDels and copy number variation (CNV). In addition, we found haploblocks with greatly increased SNP densities. We analysed these SVs and gene presence/absence polymorphisms in detail and manually validated them to distinguish true SVs from artefacts that were due to mis-assembly or annotation problems.

## Results

### Two-way comparison of Chinese Spring and ‘CH Campala *Lr22a*’ allows identification of large structural variations

Previously, 10,344 sequence scaffolds were produced from isolated chromosome 2D of ‘CH Campala *Lr22a*’ by using Chicago long-range linkage [21, 32]. In the resulting ‘CH Campala *Lr22a*’ pseudomolecule, 7,617 scaffolds were anchored, of which 7,314 were smaller than 5 kb and 90 scaffolds were larger than 1 Mb in size. The pseudomolecule had a scaffold N50 of 8.78 Mb (N90 of 1.89 Mb) and represented 98.92% of the total length of the initial assembly. To identify large InDels, we compared the Chinese Spring and ‘CH Campala *Lr22a*’ pseudomolecules in windows of 10 Mb and performed dot plots. Here, we focused only on InDels larger than 100 kb because such SVs could not be identified with previous whole-genome assemblies. In total, we found 26 putative InDels which were manually validated by evaluating the upstream and downstream sequences for the presence of ‘Ns’ at the breakpoints. If ‘Ns’ were found exactly at the breakpoints on both sides of an InDel, we considered it a false positive that was most likely due to the incorrect placement of a scaffolds in either of the pseudomolecules. Based on this criterion, we discarded 22 of the 26 candidate InDels. Three of the remaining four InDels showed good sequence quality and had clear breakpoints at both ends with no ‘Ns’. These true InDels were 285 kb, 494 kb and 765 kb in size. An additional 677 kb InDel had a clear break only at one end and ‘Ns’ on the other end. Interestingly, three of the four large InDels showed CNV for nucleotide binding site - leucine-rich repeat (NLR) genes.

Various molecular mechanisms have been described that lead to SVs. For example, unequal crossing over can occur in regions with extensive sequence similarity. On the other hand, non-homologous end-joining (NHEJ) is associated with DNA repair in regions with no or low sequence similarity. Other causes of SVs include double-strand break (DSB) repair via single-strand annealing or synthesis-dependent strand annealing mechanisms, transposable element (TEs)-mediated mechanisms and replication-error mechanisms [33-36]. These mechanisms have been well studied in humans, but in plants our understanding of the molecular causes of SVs is limited [33]. To decipher the mechanistic bases of the observed SVs, the sequence of the SV as well as their flanking regions were analyzed to identify signature sequence motifs that could point to the underlying molecular mechanism (e.g. DNA repair, recombination or replication associated mechanisms).

### Unequal crossing over is the likely cause of a 285 kb deletion in Chinese Spring

The terminal 10 Mb of the short chromosome arm revealed an InDel of 285 kb (Fig. 1a). We extracted and checked the sequences 5 kb upstream and downstream of the breakpoints for the presence of TEs or genes (or any kind of repeated sequence) that could have served as a template for unequal crossing over. Unequal crossing over occurs frequently at repeated sequences that are in the same orientation, leading to duplications or deletions of the region between the two repeats [37]. Indeed, the breakpoints of the InDel contained two NLR genes that shared 96-98% nucleotide identity in ‘CH Campala *Lr22a*’ In contrast, Chinese Spring only carried a single NLR copy (Fig. 1). Thus, it is possible that an unequal crossing over between the two genes occurred in an ancestor of Chinese Spring, leading to the loss of the 285 kb segment between the two NLRs.

**Fig. 1.**
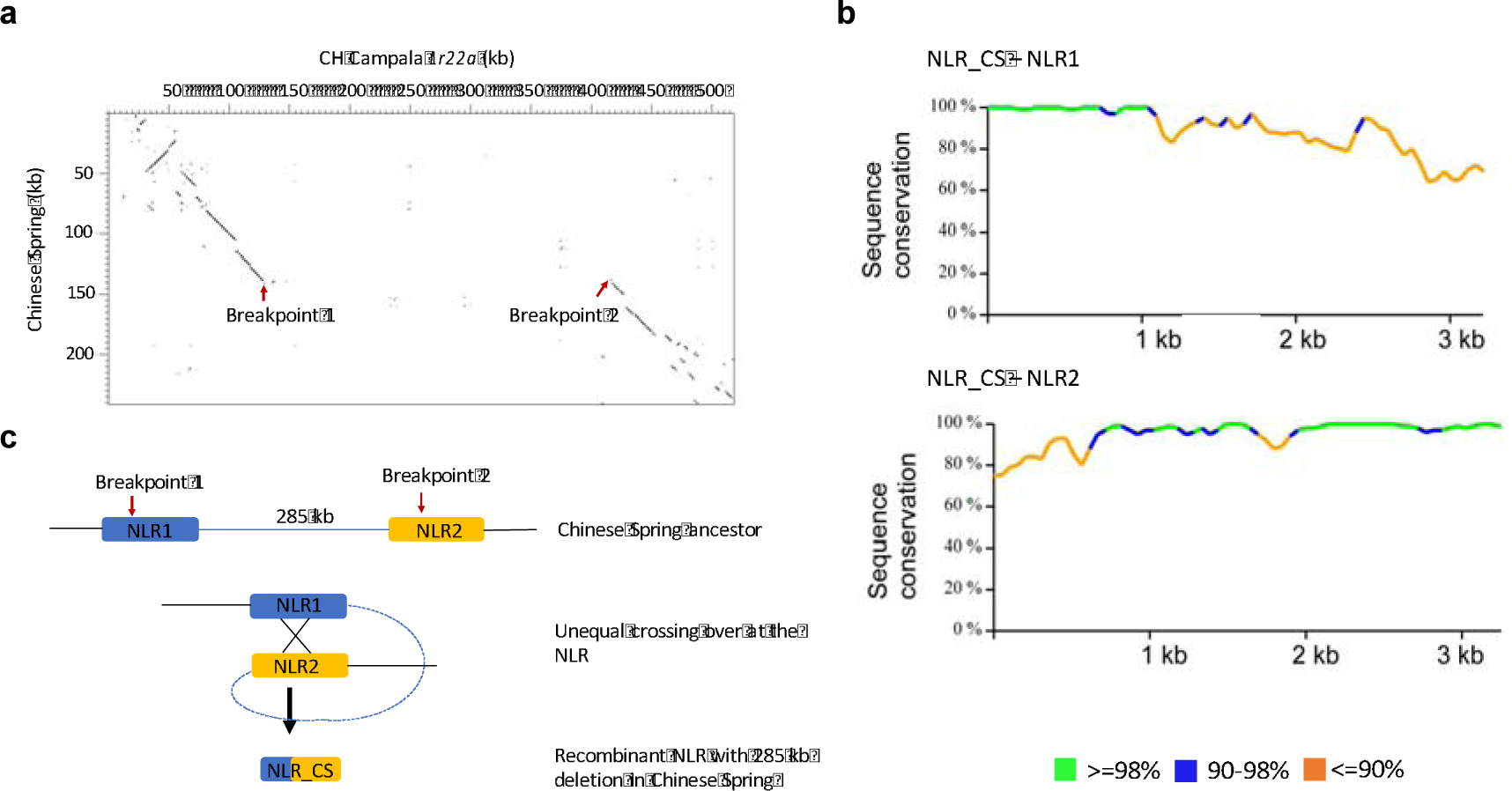
Unequal crossing over resulted in a 285 kb deletion in Chinese Spring. **a** Dot plot of a 525 kb segment from ‘CH Campala *Lr22a*’ against the corresponding 280 kb segment from Chinese Spring. The breakpoints of the 285 kb deletion are indicated by red arrows. **b** Pairwise alignment of the Chinese Spring NLR with the two ‘CH Campala *Lr22a*’ NLRs shows putative recombination breakpoints that led to the formation of the Chinese Spring NLR. **c** Proposed model for molecular events that led to a 285 kb deletion in Chinese Spring. An unequal crossing over event involving two NLR genes (shown in blue and orange) led to the formation of the recombinant NLR in Chinese Spring which shares sequence homology with NLR1 (blue) and NLR2 (yellow) and a deletion of the intervening 285 kb sequence.

In order to test this hypothesis, we further analysed the NLRs that were present at the breakpoint of ‘CH Campala *Lr22a*’ and Chinese Spring. Interestingly, the 5’ region of the Chinese Spring gene showed greater sequence similarity to NLR1 of ‘CH Campala *Lr22a*’ whereas the 3’ region was more similar to NLR2 (Fig. 1b). This suggests that these NLRs (NLR1 and NLR2) were indeed the template for an unequal crossing over in an ancestor of Chinese Spring (Fig. 1c). The corresponding 285 kb segment in ‘CH Campala *Lr22a*’ only contained repetitive sequences and did not carry any genes.

### Double-strand break repair likely mediated a large 494 kb deletion

The second SV was located on a ‘CH Campala *Lr22a*’ scaffold of 6.6 Mb in size (Fig. 2a). We could precisely identify the breakpoints based on the sequence alignment of the two wheat lines. Unlike the case described above, the upstream and downstream sequences contained no obvious sequence template or a typical TE insertion or excision pattern [34] that could have led to a large deletion by unequal crossing over. However, the breakpoints of the InDel contained typical signatures of DSB repair. In ‘CH Campala *Lr22a*’ the nucleotide triplet ‘CGA’ was repeated at both ends of the breakpoint whereas Chinese Spring had only one copy of the ‘CGA’ triplet (Fig. 2b). The proposed model for this 494 kb deletion is that it was caused through a DSB that was repaired by the single-strand annealing pathway (Fig. 2c). After the DSB that could have occurred anywhere on the 494 kb segment in Chinese Spring, 3’ overhangs were produced by exonucleases. Various studies in yeast have shown that these overhangs can be several kb in size [38-40] and due to high conservation of DSB repair pathways [41], it is expected that plants would have a similar DSB repair mechanism. In the case described here, we propose that exonucleases produced overhangs of 200-250 kb, which were then repaired by non-conservative homologous recombination repair (HRR). For this, the generated 3’ overhangs annealed in a place of complementary micro-homology, which are typically a few bp in size (‘CGA’ triplet in this case) [42]. After annealing of the matching motifs, second strand synthesis took place and the overhangs were removed, leading to the observed deletion of the 494 kb sequence in Chinese Spring (Fig. 2c). This 494 kb segment in ‘CH Campala *Lr22a*’ contained eight genes coding for an NLR, a serine/threonine protein kinase, a zinc finger-containing protein, a transferase, two cytochrome P450s and two proteins of unknown function. BLAST analysis of these eight genes against all the Chinese Spring chromosmes revealed that the homoeologous segments on the A and B genomes were retained. In other words, the deletion of these eight genes might not have led to a deleterious effect because the homoeologous gene copies on the other two sub-genomes compensate for the D-genome deletion. It has been reported that polyploid species show a higher plasticity compared to diploid species and that they are able to buffer large insertions and deletions on one particular sub-genome [43].

**Fig. 2.**
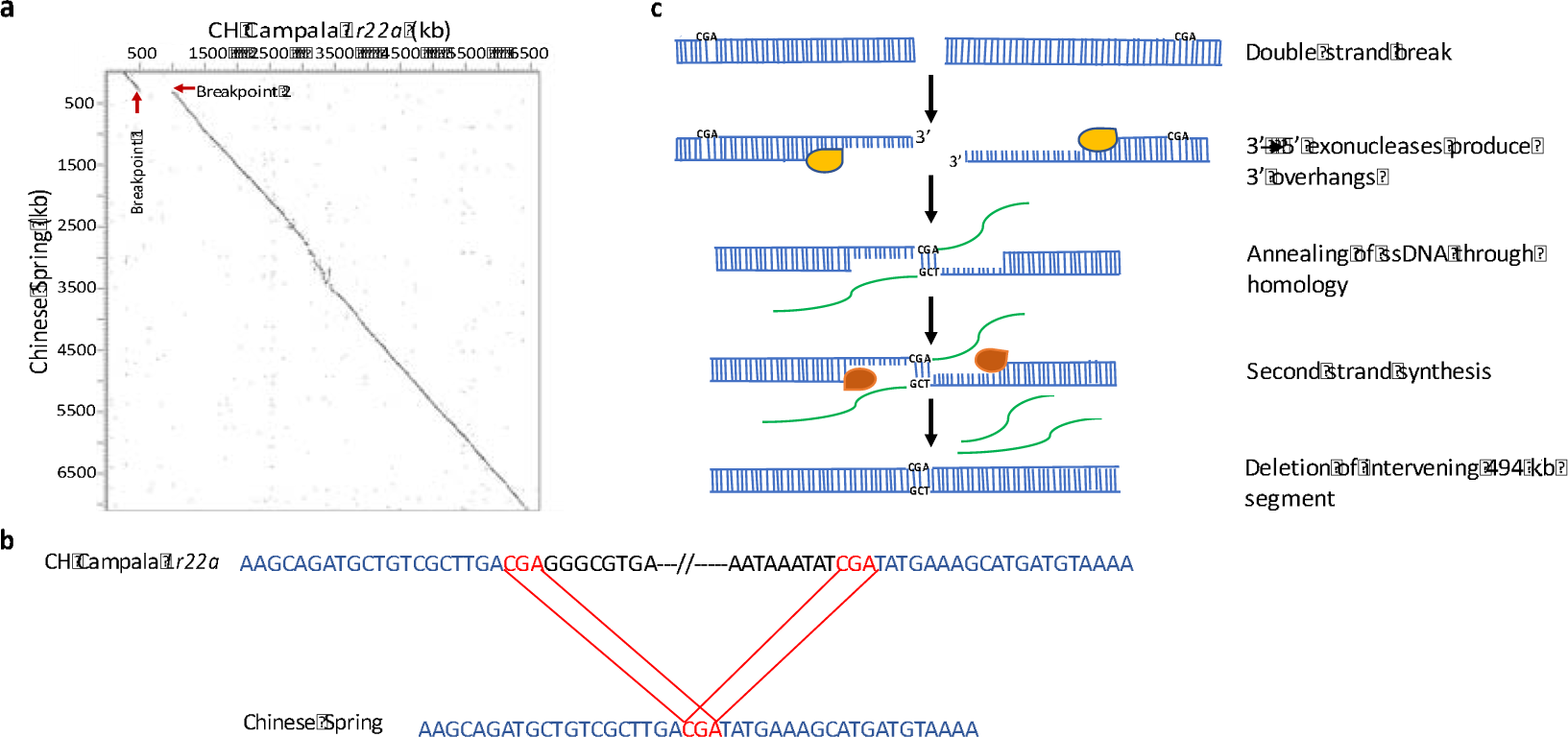
Double-strand break repair is responsible for the deletion of a 494 kb segment in Chinese Spring. **a** Dot plot of a 6.6 Mb scaffold of ‘CH Campala *Lr22a*’ against the corresponding segment from Chinese Spring. The breakpoints are indicated by red arrows. **b** Presence of DSB signatures (‘CGA’ triplet, red) with two copies in ‘CH Campala *Lr22a*’ and one in Chinese Spring. The conserved sequence is shown in blue and the 494 kb sequence that is deleted in Chinese Spring but present in ‘CH Campala *Lr22a*’ is indicated in black. **c** The proposed model for the deletion of the 494 kb segment in Chinese Spring through DSB repair by non-conservative homologous recombination repair (HRR) where the yellow enzyme is the exonuclease, green strands are the overhangs and the orange color represents the replication complex.

### Large diverse haploblocks indicate recurrent gene flow from distant relatives

Comparison of SNP density across the chromosome revealed three large regions (haploblocks *b* and c) with increased SNP density compared to the rest of the chromosome (Fig 3a). Two of the regions were located on the short arm of the chromosome whereas the largest diverse haploblock of ~48 Mb was located towards the telomeric end of the long chromosome arm. While the SNP density along most of the chromosome was in the range ~27 SNPs/Mb (Fig. 3a) the three diverse haploblocks had SNP densities of 2,500 – 4,500 SNPs/Mb. The actual number of polymorphisms might be even higher because SNP calling might not have been possible in many parts of the haploblocks because of the high sequence divergence.

**Fig. 3.**
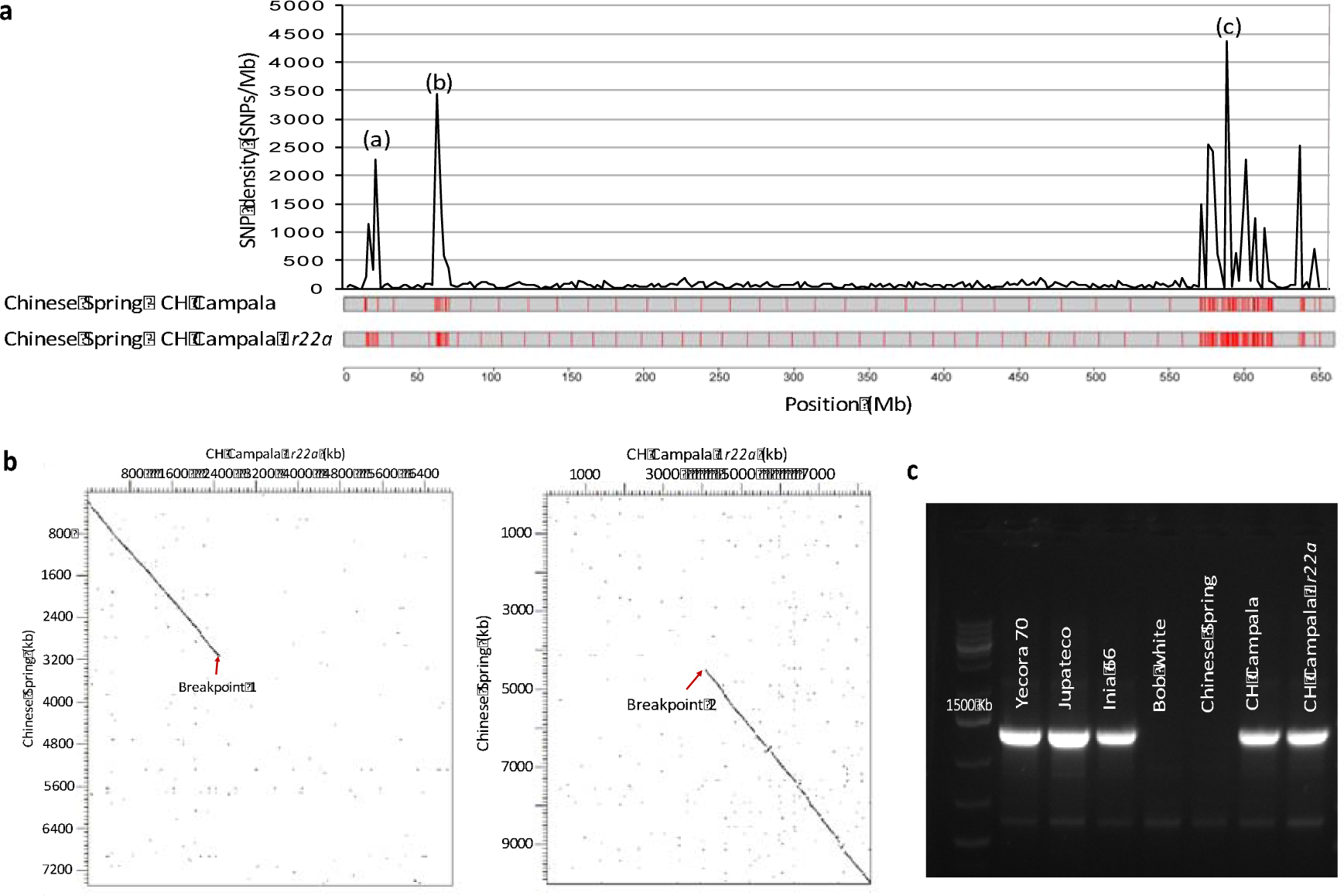
Identification of three diverse haploblocks with increased SNP density. **a** Single nucleotide polymorphism (SNP) density between Chinese Spring and ‘CH Campala *Lr22a*’ in a sliding windows of 2.5 Mb. The numbers refer to the position in Mb along the chromosome 2D of the Chinese Spring. The three diverse haploblocks are indicated with letters (a), (b) and (c). **b** Dot plot of Chinese Spring and ‘CH Campala *Lr22a*’ showing the left and right breakpoints of the large haploblock *c.* The sequence adjacent to the haploblock shows a high degree of sequence conservation in intergenic regions whereas the sequence similarity was very low in the haploblock region. **c** PCR amplification using an introgression-specific primer pair designed on the left breakpoint of the ‘CH Campala *Lr22a*’ introgression. Jupateco, Yecora 70 and Inia 66 are CIMMYT wheat cultivars. Inia 66 is in the pedigree of ‘CH Campala *Lr22a*’.

The first haploblock (haploblock *a*) at the distal end of the short chromosome arm contains the *Lr22a* leaf rust resistance gene that was introduced into hexaploid wheat through an artificial hybridization between a tetraploid wheat line and an *Ae. tauschii* accession [44]. There are two genetically distant lineages of *Ae. tauschii.* The D-genome of hexaploid wheat was most likely contributed by an *Ae. tauschii* population belonging to lineage 2 [45], whereas the donor of *Lr22a* (*Ae. tauschii* accession RL 5271) belongs to the genetically diverse lineage 1 [46]. The size of the *Lr22a* introgression was subsequently reduced through several rounds of backcrossing with hexaploid wheat and the remaining *Lr22a*-containing segment was bred into elite wheat lines including ‘CH Campala *Lr22a*’ to increase resistance against the fungal leaf rust disease [31]. Based on the SNP density, we were able to estimate the size of the remaining, introgressed *Ae. tauschii* segment to ~8 Mb. The original donor of the other two haploblocks (haploblocks *b* and *c*) could not be traced back and they might be the result of natural gene flow or artificial hybridization. Mapping of independently generated short-read sequences from ‘CH Campala’, the recurrent parent that was used to produce the near isogenic line ‘CH Campala *Lr22a*’, showed that the same haploblocks were also present in ‘CH Campala’ (Fig. 3a), indicating that these segments were not co-introduced along with the *Lr22a* segment from RL 5271. In particular, the presence of the large continuous haploblock *c* on the long chromosome arm was intriguing. Dot plots allowed us to identify the exact breakpoints of the haploblock (Fig. 3b). While there was high sequence homology in both flanking regions, sequence identity in the intergenic regions broke down inside the haploblock (Fig. 3b). In contrast, dot plots with haploblocks *a* and *b* revealed a good level of collinearity between Chinese Spring and ‘CH Campala *Lr22a*’ in intergenic regions despite the increased SNP density (Additional file 1: Figure S1), indicating that haploblock *c* is the most diverse. Comparison to the recently generated high-quality genome assembly of *Ae. tauschii* accession AL8/78 [47], an accession that is closely related to the wheat D-genome and that belongs to lineage 2, suggests that haploblock *c* represents an interstitial introgression into ‘CH Campala *Lr22a*’ (Additional file 2: Figure S2). In Chinese Spring, 723 genes were located in this haploblock, whereas ‘CH Campala *Lr22a*’ contained 681 genes in this region. The genic sequences in the haploblock region showed a nucleotide sequence identity of 78-100% compared to 99-100% for the genes outside the haploblock. We also observed three inversions of —1.48 Mb, ~422 kb and ~418 kb in the haploblock *c* where the gene order was reversed.

To track the possible origin of this introgression, we developed an introgression-specific PCR probe based on the sequence of the left breakpoint in ‘CH Campala *Lr22a*’. The marker amplified in several wheat cultivars that were developed by the International Wheat and Maize Improvement Center (CIMMYT) (Fig. 3c). Among them is Inia-66, which is in the pedigree of ‘CH Campala *Lr22a*’ [48]. These results indicate that the particular segment in ‘CH Campala *Lr22a*’ might have been introgressed via a CIMMYT cultivar.

### Chromos ome-wi de comparison of NLR genes reveals extensive copy number variation in certain NLR families

Regions harboring NLR genes have been reported to be fast evolving to keep up in the arms-race with pathogens [49]. Interestingly, three of the four large InDels identified created CNV for NLR genes. We were therefore interested in the evolutionary dynamics of chromosomal regions harboring NLR genes. For chromosome 2D, a total of 161 NLRs were annotated in the wheat line ‘CH Campala *Lr22a*’ and 158 NLRs for Chinese Spring. The NLRs annotated in the two wheat genotypes showed a high tendency of clustering and they were mostly located in the telomeric regions (Fig. 4a), as it is typically found for this gene class [25].

**Fig. 4.**
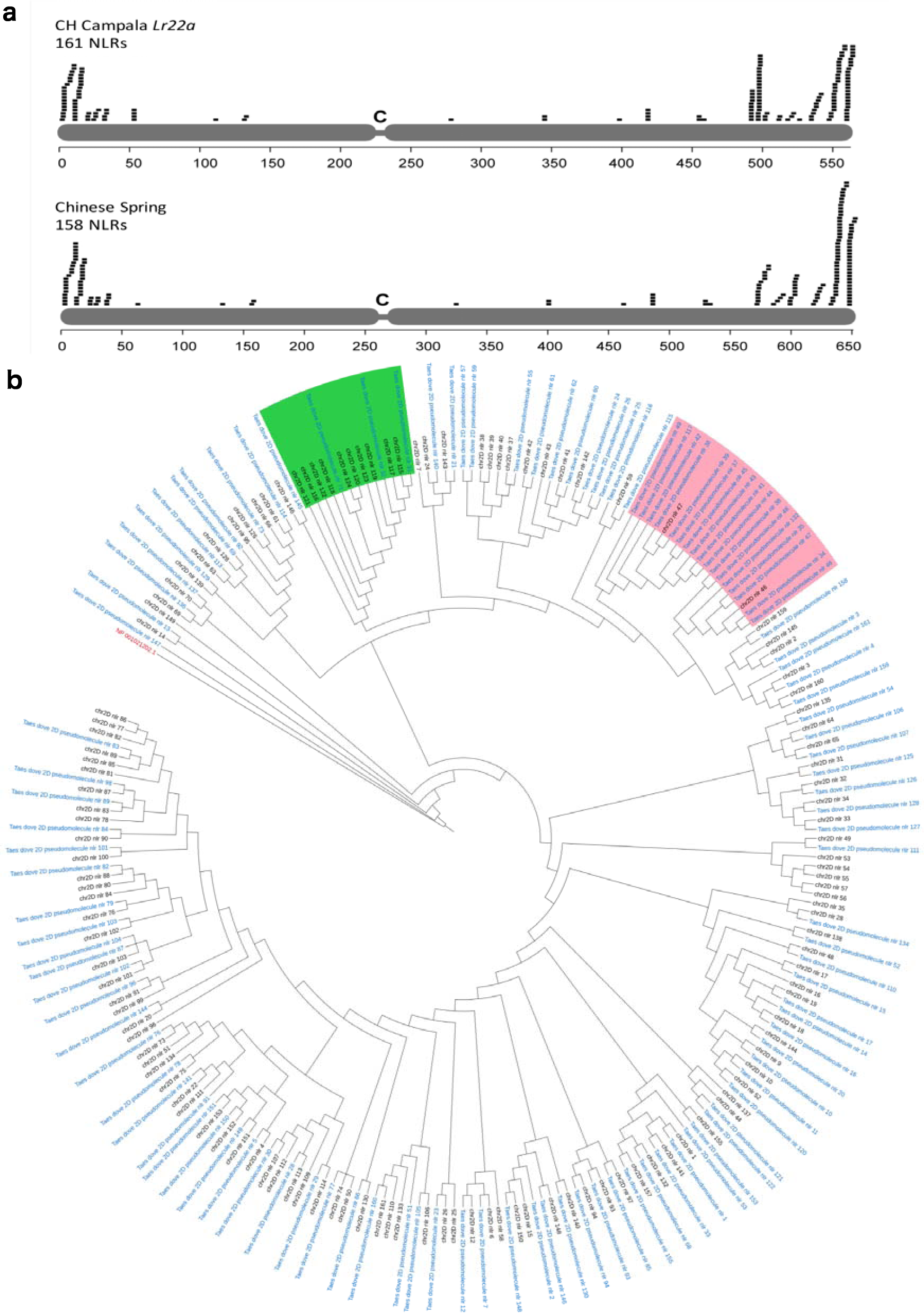
Distribution of predicted NLR genes on chromosome 2D. **a** The x-axis indicates the position in Mb. Note that the scales differ between ‘CH Campala *Lr22a*’ and Chinese Spring, because the sequence assembly of ‘CH Campala *Lr22a*’ is shorter than that of Chinese Spring. **b** Phylogenetic tree where blue labels ‘Taes dove 2D pseudomolecule nlr’ represent the ‘CH Campala *Lr22a*’ NLRs and black labels ‘chr2D nlr’ represent the Chinese Spring NLRs. The two highlighted regions in green and pink represent chromosomal segments with high copy number variation that are discussed in the text.

For ‘CH Campala *Lr22a*’. we found that 62 NLR genes resided in seven gene clusters which comprise 38.5% of the total annotated NLRs. The largest cluster contained 19 NLR genes. In Chinese Spring, we found that 71 NLR genes resided in ten clusters which comprise 44.9% of the total annotated NLRs and the largest cluster contained 21 NLRs. A phylogenetic tree revealed that most NLR genes from Chinese Spring had one ortholog in ‘CH Campala *Lr22a*’ (Fig. 4b). On the other hand, we also observed copy number variation for certain regions. Two regions, CNV1 and CNV2, were of particular interest because there was an extensive variation in the NLR copy number between Chinese Spring and ‘CH Campala *Lr22a*’ (Fig. 4b). In the CNV1 region, ‘CH Campala *Lr22a*’ had sixteen NLR genes annotated in a 786 kb region. The corresponding region in Chinese Spring contained only two NLRs in a 21 kb interval (Fig. 5a). There was a high degree of gene collinearity flanking the NLR cluster (Fig. 5a). The two NLR copies in Chinese Spring (NLR46 and NLR47) showed 44% sequence identity at the protein level, indicating that they might have arisen from a very ancient gene duplication. The low sequence identity of NLR46 and NLR47 allowed to assign each of the ‘CH Campala *Lr22a*’ NLRs to one of the two Chinese Spring copies. This revealed a random pattern, which might be explained by complex duplication and rearrangement events (Fig. 5a). The CNV1 region locates to the diverse haploblock *c*, which might explain the extent of the CNV found in this region.

**Fig. 5.**
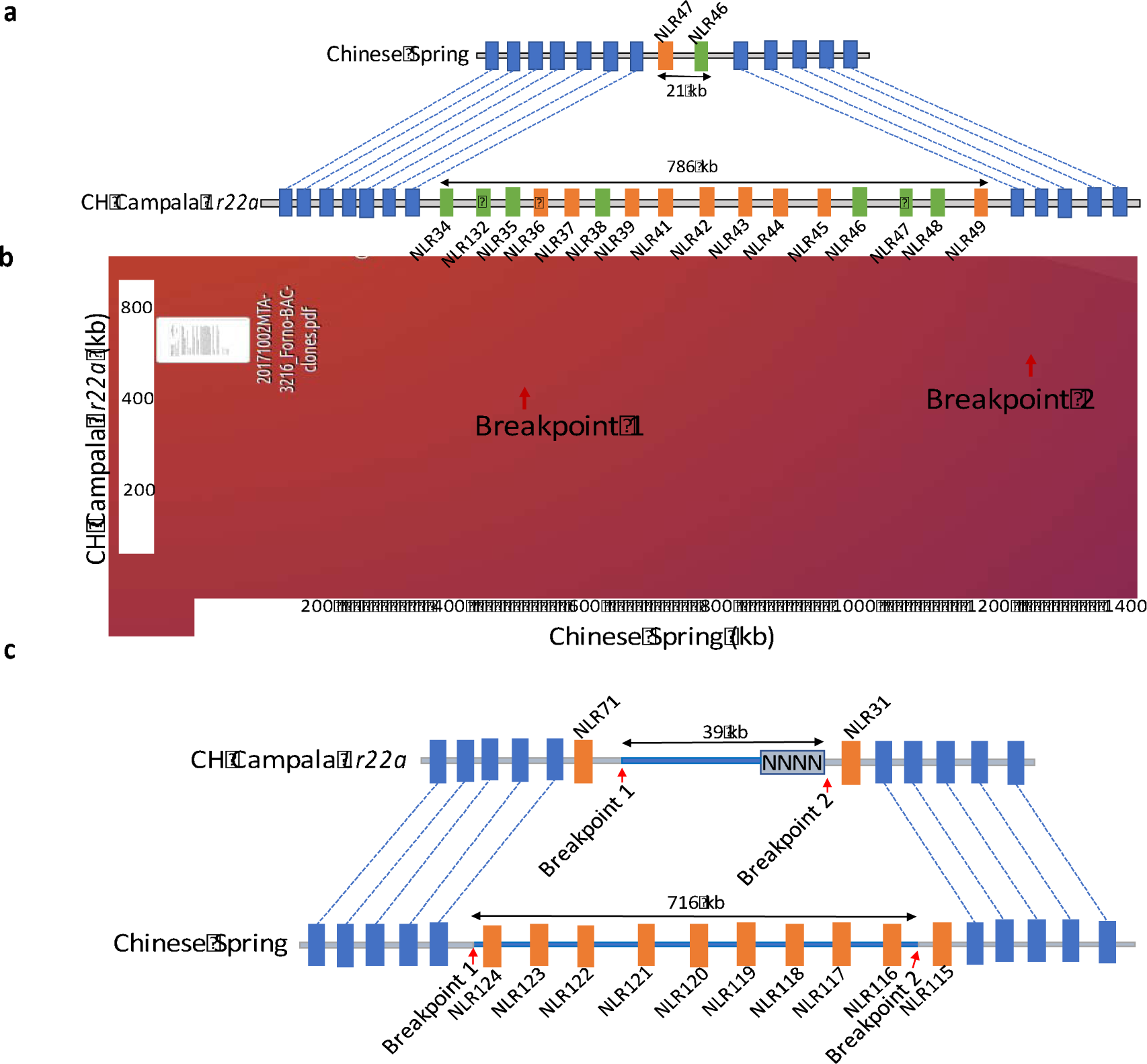
NLR copy number variation. **a** In the CNV1 region we found 16 NLRs in ‘CH Campala *Lr22a*’ annotated in a 786 kb region. Pseudogenes are marked with J. Chinese Spring has only two NLRs in a 21 kb segment. **b** NLR gene expansion in Chinese Spring. Dot plot of the CNV region between Chinese Spring and ‘CH Campala *Lr22a*’ **c** Chinese Spring had 21 NLRs compared to 14 in ‘CH Campala *Lr22a*’ which are shown in orange and the collinear genes in the flanking region are shown in blue.

The CNV2 region affected a segment of ten paralogous NLR genes situated in a 716 kb region in Chinese Spring. In ‘CH Campala *Lr22a*’ there was a 677 kb deletion that affected all but two of the NLRs. For this CNV region we could identify a clear breakpoint at one end whereas the other end had a sequence gap (Fig. 5b and 5c).

## Discussion

### Molecular mechanisms of structural variations

Different genotypes within a plant species can show tremendous genetic diversity. Beside SNPs, SVs have been identified as a major contributor to phenotypic variation in plants, which is why an understanding of large SVs is of importance for breeding [50]. For example, the durable fungal stem rust resistance gene *Sr2* of wheat was localized to a region on chromosome 3B that showed extensive structural rearrangements between the Sr2-carrying wheat cultivar Hope and the susceptible Chinese Spring on an 867 kb chromosome segment [51]. How this structural rearrangement affects the Sr2-mediated stem rust resistance is not yet understood. Similarly, large deletions comprising multiple tandemly duplicated transcription factor genes at the *Frost resistance-2* locus are associated with reduced frost tolerance in wheat [52]. While short-read sequencing allowed a comprehensive assessment of genome-wide SNPs distributions in cereals [53, 54] the identification of SVs, particularly large InDels, has been challenging due to technical limitations. In wheat, the lack of high-quality chromosome assemblies from multiple genotypes has prevented such comparisons so far. Even for other cereal crop species like rice, maize, barley and sorghum there are no or only very few high-quality *de novo* assemblies available beside the reference genotypes [22, 55-57]. Here, we compared two high-quality sequence assemblies of bread wheat chromosome 2D that were highly contiguous over megabases, which allowed us to focus on InDels of several hundred kb in size. In total, we found that around 0.3% of the chromosome was affected by the four large InDels. Based on these numbers, we estimate that a comparison of any two wheat genotypes would reveal around 30 large InDels affecting ~15 Mb accross the entire D sub-genome. Not surprisingly, the number of small InDels is much higher than larger structural rearrangements. For example, a comparison of the B73 maize reference assembly to optical maps generated from the two maize inbred lines Ki11 and W22 revealed around 3,400 insertions and deletions between two maize lines with an average InDel size of 20 kb [17]. A re-sequencing study in rice revealed a total of 13,045 insertions and 15,151 deletions in the size range of 10-1,000 bp [58]. Large InDels affected multiple genes and can therefore have a deleterious effect, particularly in diploid species.

Unequal crossing over and DSB repair were identified as the molecular mechanisms responsible for large InDels in our study. Analyses in Brachypodium revealed that DSB repair is the most common mechanism for structural rearrangements [34, 59]. The error prone DSB repair leads to insertions, deletions or rearrangements in the genome. In our comparative analysis, we found a large deletion of 494 kb in Chinese Spring where DSB repair via single strand annealing led to the deletion of the intervening region between the conserved motifs known as DSB signatures. Similar mechanisms were identified in a comparative analysis of the two barley cultivars Barke and Morex, where DSB repair accounted for 41% of the InDel events [33]. DSB repair signatures were also found in maize where they flanked small InDels ranging from 5 bp to 175 bp [60]. Apart from DSB repair, another frequently observed mechanism for SV is unequal crossing over. We found a 285 kb deletion in Chinese Spring where the deletion was a result of an improper alignment of two highly similar NLR genes that served as a template for unequal crossing over. Unequal crossing over has been shown to be one of the main driving forces for genome evolution and has been reported to occur in various disease resistance gene families where they result in novel specificities and haplotypes [37]. For example, unequal crossing over between homologs in the maize rust resistance locus *Rp1* led to the formation of recombinant genes with diverse resistance specificities [61, 62]. In soybean, unequal crossing over at the *RPS* locus was associated with loss of resistance to Phytophthora due to the deletion of a NLR-like (*NBSRps4/6*) sequence [63].

### Identification of diverse haploblocks – implications for wheat D-genome evolution

In addition to SVs, the chromosome-scale assemblies also allowed us to assess SNP density across the entire chromosome and to identify large contiguous blocks with strong variation from the average SNP density. This revealed the presence of three large haploblocks that showed a much higher SNP density compared to the rest of the chromosome. One of these haploblocks (haploblock *a*) could be traced back to an artificial introgression that carries the adult plant leaf rust resistance gene *Lr22a* [32, 64]. *Lr22a* was introgressed into hexaploid wheat by artificially hybridizing the tetraploid wheat line tetra-Canthatch with the diploid *Ae. tauschii* accession RL 5271 [44]. The crossing of tetraploid wheat with diverse *Ae. tauschii* accessions results in so called synthetic wheat. This is a widely explored strategy in breeding to compensate for the loss of diversity in hexaploid wheat that went along with domestication and modern breeding [65-67]. After this initial cross, the resulting synthetic hexaploid wheat line was backcrossed six times with the historically important North American wheat cultivar Thatcher, which resulted in the Lr22a-containing backcross line ‘Thatcher *Lr22a*’ (RL 6044). This backcross line then served as the donor to transfer *Lr22a* into elite wheat cultivars including the Canadian wheat cultivar ‘AC Minto’ and the Swiss spring wheat line ‘CH Campala *Lr22a*’ [31, 64]. The SNP density analysis allowed us now to precisely determine the size of the remaining RL 5271 segment after a limited number of crosses. We did not find evidence for co-introduction of additional segments from the original *Ae. tauschii* donor along chromosome 2D. More interestingly, two additional diverse haploblocks (haploblocks *b* and c) of almost 9 Mb and 48 Mb were identified towards the telomeric end of the short and long chromosome arms, respectively. It has been reported that the wheat D genome was most likely contributed by an *Ae. tauschii* population from a region close to the southern or southwestern Caspian Sea. This accession belonged to one of two genetically distinct sublineages within the *Ae. tauschii* gene pool (sublineage 2) [45]. However, it has been found that gene flow from *Ae. tauschii* accessions belonging to the genetically distant sublineage 1 occurred after the formation of hexaploid wheat, which might explain the presence of contiguous haploblocks with increased diversity. Interestingly, Wang et al. (2013) [45] identified a putative introgression of *Ae. tauschii* sublineage 1 on the telomeric end of chromosome arm 2DL in hexaploid wheat, which might be identical to the diverse haploblock *c* identified in our study. Alternatively, these diverse haploblocks might stem from an alien introgression from another grass species. Interspecies hybridizations are a common method in wheat breeding to transfer specific traits from wild and domesticated grasses into wheat [68]. In contrast to the naturally occurring gene flow from *Ae. tauschii*, the vast majority of these alien introgressions were artificially produced and require in-vivo culture techniques like embryo rescue. The length of the haploblock *c* was surprising because the size of haploblocks is expected to be negatively correlated with recombination rates [69]. Since the haploblock *c* located to the highly-recombining telomeric end of the chromosome, we would expect that its size decreases over time. One explanation for conservation of this haploblock could be that its presence suppresses recombination in this area. In contrast to haploblocks *a* and *b*, we observed a breakdown of sequence homology in intergenic regions in haploblock *c*. On the other hand, the gene order was largely collinear in haploblock *c*, which should be sufficient for recombination in this chromosome segment. A second explanation is that this haploblock *c* might be widely present in the wheat gene pool or in particular breeding programs. For example, PCR analysis revealed that the haploblock *c* was present in multiple CIMMYT wheat lines. This would allow recombination in the haploblock without decreasing its size. In summary, a considerable fraction of the chromosome (10%) was made up of haploblocks with a much greater diversity than the rest of the chromosome. This highlights the importance of natural gene flow and artificial hybridization as sources for diversity in cereal breeding.

## Conclusions

This study provides the first comparison of two pseudomolecules based on high-quality *de novo* chromosome assemblies. The megabase-sized scaffolds allowed us to focus particularly on InDels of several hundred kb in size. Our analysis revealed that around 0.3% of the chromosome was affected by large InDels between the two wheat lines. Our study also revealed that careful manual validation is required in order not to overestimate the frequency of InDels. In particular, 84% of the InDels that were initially identified were removed after manual curation because they were most likely due to assembly and annotation artefacts. It is conceivable that previous comparative analyses in wheat that were based on short-read resequencing alone could not account for these problems. We therefore highlight the importance of manual data validation in future wheat pan-genome projects.

## Methods

### ‘CH Campala *Lr22a*’ pseudomolecule assembly

The initial sequence assembly provided by Dovetail Genomics consisted of 10,344 sequence scaffolds (hereafter referred to as Dovetail scaffolds) with and average size of 54.8 kb and an N50 of 9.758 Mb [32]. To anchor these scaffolds, segments of the scaffolds were used in BLASTN searches against the Chinese Spring chromosome [25]. Dovetail scaffolds shorter than 10 kb were used in their entirety for the BLASTN search. For Dovetail scaffolds between 10 and 200 kb, a 1 kb segment every 30 kb was used for the BLASTN search. For Dovetail scaffolds larger than 200 kb, a 1 kb segment every 100 kb was used for BLASTN search. For each Dovetail scaffold, it was then determined where the majority of BLAST hits were located in Chinese Spring 2D. Based on this information, Dovetail scaffolds were ordered.

After sequence scaffolds were assembled into a first version of a pseudomolecule, we searched for large-scale breaks in gene collinearity when compared to Chinese Spring chromosome 2D. Here, we focused on blocks of BLASTN hits that mapped to completely different regions of the genome. If the end of a non-collinear block coincided with the end of a Dovetail scaffold, this was interpreted as an assembly artefact. The approximate location of the mis-assembly was identified and the respective Dovetail scaffold was then split into segments. We identified ten putatively chimeric Dovetail scaffolds with assembly errors. These were split into 24 segments (some Dovetail scaffolds contained multiple mis-assemblies) which were then anchored individually to Chinese Spring chromosome 2D.

A total of 7,617 Dovetail scaffolds were integrated to the final pseudomolecule of 563 Mb, representing 73% of all Dovetail scaffolds and 98.92% of the total length of the Dovetail assembly. The integrated 7,617 Dovetail scaffolds have an N50 of 8.78 Mb and an N90 of 1.89 Mb. The scaffold N50 of 8.78 Mb is slightly lower than the N50 of the original assembly obtained from Dovetail Genomics, which is due to the splitting of chimeric scaffolds.

### NLR annotation and phylogenetic tree

NLR loci on the ‘CH Campala *Lr22a*’ pseudomolecule was annotated using NLR-Annotator [73]. The initial fragmentation step of NLR-Annotator was performed generating 20 kb fragments that overlap by 5 kb. Multiple alignments of NB-ARC associated amino acid motifs were generated using NLR-Annotator (output option −a). Multiple alignment files were concatenated and a comparative phylogenetic tree was generated using FastTree[74] version 2.1.7 [75].

### Identification of the SVs

We analysed SVs in the telomeric and interstitial regions and excluded the centromeric region which was ~100 Mb in size (position 190-290 Mb in Chinese Spring pseudomolecule and 150-250 Mb in ‘CH Campala *Lr22a*’ pseudomolecule). The centromeric region is extremely repetitive and gene poor and alignments were difficult. For the identification of the SVs, we segmented the Chinese Spring and ‘CH Campala *Lr22a*’ pseudomolecules in the windows of 10 Mb and performed dot plot alignments (program DOTTER) [76]. For each of the InDels observed, we analysed the sequence alignments to identify the region where the sequence similarity broke down and this region was called breakpoint. We spliced out 5 kb sequence upstream and downstream of these breakpoints and performed BLASTN search [77] against the repeat database to identify transposable elements and also against the *Brachypodium distachyon* coding sequence database [78] to identify genes in the flanking regions to understand the molecular mechanism underlying the observed SVs.

To identify NLR CNV, we compared the NLR clusters in Chinese Spring and ‘CH Campala *Lr22a*’ and identified the breakpoints as described above. The sequences upstream and downstream of breakpoints were used to identify the collinear genes using BLAST search against the annotated ‘CH Campala *Lr22a*’ and Chinese Spring genes. Putative start and stop codons of the annotated NLRs were identified based on the orthologs of these NLRs in *Brachypodium distachyon.* The coding sequence of these *Brachypodium distachyon* NLRs was taken from the *Brachypodium distachyon* coding sequence database [78] and was used for the dot plot alignment to identify the coding sequence of the Chinese Spring and ‘CH Campala *Lr22a*’ NLRs. Pseudogenes were predicted on the basis of frameshift mutations, premature stop codon or insertion of a transposable element resulting in a pseudogene.

### Haploblock analysis and validation

For the identification of the haploblock region, we mapped previously generated Illumina reads of ‘CH Campala *Lr22a*’ and ‘CH Campala’ [32] to the Chinese Spring pseudomolecule using the CLC Main Workbench 7 (Qiagen) with standard parameters. The mapped read file was later used for the variant call analysis on the CLC Main Workbench 7 (Qiagen) using standard parameters. SNP density was calculated in sliding windows of 2.5 Mb. To verify the haploblock *c* region we designed a PCR probe (forward primer-GCCACGAGCGTGGTCGTG and reverse primer-CCTTCATAGCTCCGTAGAAG) spanning the left border of the haploblock *c* of ‘CH Campala *Lr22a’*. The PCR amplification was performed in 20 μl reaction mixture containing 65 ng of genomic DNA, 1 μl of 2.5 mM dNTP’s, 1 μ! of 10 μM of each primer and 0.25 units of Sigma Taq polymerase at 60 °C annealing temperature for 35 cycles. The cycling parameters used were, pre-denaturation at 95 °C for 4 min, which was followed by 35 cycles of 95 °C for 30 s, annealing at 60 °C for 30 s, 72 °C for 2 min and a final extension at 72 °C for 10 min. The PCR products were separated on 1.0% agarose gels.

## Acknowledgements

We would like to thank Dr. Dario Fossati from Agroscope, Switzerland, Dr. Ravi Singh from CIMMYT and Prof. Jaroslav Dolezel from the Institute of Experimental Botany, Czech Republic for commenting on the possible origin of the large ‘CH Camapala *Lr22a*’ introgression.

## Availability of data and materials

The submission of ‘CH Campala *Lr22a*’ Pseudomolecule and its annotation is in progress and will be available on request.

## Author’s contributions

AKT, TW, BK and SGK conceived the study. AKT, TW and TM performed the bioinformatics analyses. PMA performed PCR experiments. BS and BBHW annotated the NLR genes. MS, ST, MF, TL and KFXM performed gene annotation in ‘CH Campala *Lr22a*’. AKT, TW, BK and SGK wrote the manuscript and all authors read and approved the final manuscript.

## Ethics approval and consent in participate

Not applicable

## References

1. The Food and Agriculture Organization of the United Nations. http://www.fao.org/faostat/en/ (2017). Accessed 12 Dec 2017.

2. Wicker T, Gundlach H, Spannagl M, Uauy C, Borrill P, Ramírez-González RH, Oliveira RD, Consortium IWGS, Mayer KFX, Paux E, Choulet F. unpublished.

3. Jordan KW, Wang S, Lun Y, Gardiner LJ, MacLachlan R, Hucl P, Wiebe K, Wong D, Forrest KL, Consortium I, et al. A haplotype map of allohexaploid wheat reveals distinct patterns of selection on homoeologous genomes. Genome Biol. 2015;16:48.

4. Avni R, Nave M, Barad O, Baruch K, Twardziok SO, Gundlach H, Hale I, Mascher M, Spannagl M, Wiebe K, et al. Wild emmer genome architecture and diversity elucidate wheat evolution and domestication. Science. 2017;357:93–7.

5. Salamini F, Ozkan H, Brandolini A, Schafer-Pregl R, Martin W. Genetics and geography of wild cereal domestication in the near east. Nat Rev Genet. 2002;3:429–41.

6. Akhunov ED, Akhunova AR, Anderson OD, Anderson JA, Blake N, Clegg MT, Coleman-Derr D, Conley EJ, Crossman CC, Deal KR, et al. Nucleotide diversity maps reveal variation in diversity among wheat genomes and chromosomes. BMC Genomics. 2010;11:702.

7. Brenchley R, Spannagl M, Pfeifer M, Barker GLA, D’Amore R, Allen AM, McKenzie N, Kramer M, Kerhornou A, Bolser D, et al. Analysis of the breadwheat genome using whole-genome shotgun sequencing. Nature. 2012;491:705–10.

8. The International Wheat Genome Sequencing Consortium. A chromosome-based draft sequence of the hexaploid bread wheat (*Triticum aestivum*) genome. Science. 2014;345:1251788.

9. Ling HQ, Zhao S, Liu D, Wang J, Sun H, Zhang C, Fan H, Li D, Dong L, Tao Y, et al. Draft genome of the wheat A-genome progenitor *Triticum urartu*. Nature. 2013;496:87–90.

10. Jia J, Zhao S, Kong X, Li Y, Zhao G, He W, Appels R, Pfeifer M, Tao Y, Zhang X, et al. *Aegilops tauschii* draft genome sequence reveals a gene repertoire for wheat adaptation. Nature. 2013;496:91–5.

11. Paux E, Sourdille P, Salse J, Saintenac C, Choulet F, Leroy P, Korol A, Michalak M, Kianian S, Spielmeyer W, et al. A physical map of the 1-gigabase bread wheat chromosome 3B. Science. 2008;322:101–04.

12. Choulet F, Alberti A, Theil S, Glover N, Barbe V, Daron J, Pingault L, Sourdille P, Couloux A, Paux E, et al. Structural and functional partitioning of bread wheat chromosome 3B. Science. 2014;345:1249721.

13. Chapman JA, Mascher M, Buluc A, Barry K, Georganas E, Session A, Strnadova V, Jenkins J, Sehgal S, Oliker L, et al. A whole-genome shotgun approach for assembling and anchoring the hexaploid bread wheat genome. Genome Biol. 2015;16:26.

14. Clavijo BJ, Venturini L, Schudoma C, Accinelli GG, Kaithakottil G, Wright J, Borrill P, Kettleborough G, Heavens D, Chapman H, et al. An improved assembly and annotation of the allohexaploid wheat genome identifies complete families of agronomic genes and provides genomic evidence for chromosomal translocations. Genome Res. 2017;27:885–96.

15. Zimin AV, Puiu D, Hall R, Kingan S, Clavijo BJ, Salzberg SL. The first near-complete assembly of the hexaploid bread wheat genome, *Triticum aestivum*. Gigascience. 2017;6:1–7.

16. Hirsch CN, Hirsch CD, Brohammer AB, Bowman MJ, Soifer I, Barad O, Shem-Tov D, Baruch K, Lu F, Hernandez AG, et al. Draft assembly of elite inbred line PH207 provides insights into genomic and transcriptome diversity in maize. Plant Cell. 2016;28:2700–14.

17. Jiao Y, Peluso P, Shi J, Liang T, Stitzer MC, Wang B, Campbell MS, Stein JC, Wei X, Chin CS, et al. Improved maize reference genome with single-molecule technologies. Nature. 2017;546:524–7.

18. Schmidt MHW, Vogel A, Denton AK, Istace B, Wormit A, van de Geest H, Bolger ME, Alseekh S, Mass J, Pfaff C, et al. De novo assembly of a new *Solanum pennellii* accession using nanopore sequencing. Plant Cell. 2017;29:2336–48.

19. Lieberman-Aiden E, van Berkum NL, Williams L, Imakaev M, Ragoczy T, Telling A, Amit I, Lajoie BR, Sabo PJ, Dorschner MO, et al. Comprehensive mapping of long-range interactions reveals folding principles of the human genome. Science. 2009;326:289–93.

20. van Berkum NL, Lieberman-Aiden E, Williams L, Imakaev M, Gnirke A, Mirny LA, Dekker J, Lander ES. Hi-C: a method to study the three-dimensional architecture of genomes. J Vis Exp. 2010;39:1869.

21. Putnam NH, O’Connell BL, Stites JC, Rice BJ, Blanchette M, Calef R, Troll CJ, Fields A, Hartley PD, Sugnet CW, et al. Chromosome-scale shotgun assembly using an in vitro method for long-range linkage. Genome Res. 2016;26:342–50.

22. Mascher M, Gundlach H, Himmelbach A, Beier S, Twardziok SO, Wicker T, Radchuk V, Dockter C, Hedley PE, Russell J, et al. A chromosome conformation capture ordered sequence of the barley genome. Nature. 2017;544:427–33.

23. Jarvis DE, Ho YS, Lightfoot DJ, Schmockel SM, Li B, Borm TJ, Ohyanagi H, Mineta K, Michell CT, Saber N, et al. The genome of *Chenopodium quinoa*. Nature. 2017;542:307–12.

24. Moll KM, Zhou P, Ramaraj T, Fajardo D, Devitt NP, Sadowsky MJ, Stupar RM, Tiffin P, Miller JR, Young ND, et al. Strategies for optimizing BioNano and Dovetail explored through a second reference quality assembly for the legume model, Medicago truncatula. BMC Genomics. 2017;18:578.

25. The International Wheat Genome Sequencing Consortium. unpublishedf.

26. Endo TR, Gill BS. The deletion stocks of common wheat. J Hered. 1996;87:295–307.

27. Sears ER, Sears LMS. The Telocentric Chromosomes of Common Wheat. In: S. Ramanujam, editor. Proceedings of the 5th International Wheat Genetics Symposium, New Delhi; 1978. p. 389–407.

28. Alkan C, Coe BP, Eichler EE. Genome structural variation discovery and genotyping. Nat Rev Genet. 2011;12:363–76.

29. Liu M, Stiller J, Holusova K, Vrana J, Liu D, Dolezel J, Liu C: Chromosome-specific sequencing reveals an extensive dispensable genome component in wheat. Sci Rep. 2016;6:36398.

30. Montenegro JD, Golicz AA, Bayer PE, Hurgobin B, Lee H, Chan CK, Visendi P, Lai K, Dolezel J, Batley J, Edwards D. The pangenome of hexaploid bread wheat. Plant J. 2017;90:1007–13.

31. Moullet O, Schori A. Maintaining the efficiency of MAS method in cereals while reducing the costs. J Plant Breed Genet. 2014;2:97–100.

32. Thind AK, Wicker T, Simkova H, Fossati D, Moullet O, Brabant C, Vrana J, Dolezel J, Krattinger SG. Rapid cloning of genes in hexaploid wheat using cultivar-specific long-range chromosome assembly. Nat Biotechnol. 2017;35:793–6.

33. Munoz-Amatriain M, Eichten SR, Wicker T, Richmond TA, Mascher M, Steuernagel B, Scholz U, Ariyadasa R, Spannagl M, Nussbaumer T, et al. Distribution, functional impact, and origin mechanisms of copy number variation in the barley genome. Genome Biol. 2013;14:R58.

34. Wicker T, Buchmann JP, Keller B. Patching gaps in plant genomes results in gene movement and erosion of colinearity. Genome Res. 2010;20:1229–37.

35. Wicker T, Yu YS, Haberer G, Mayer KFX, Marri PR, Steve RW, Chen MS, Zuccolo A, Panaud O, Wing RA, Roffler S. DNA transposon activity is associated with increased mutation rates in genes of rice and other grasses. Nature Commun. 2016;7:12790.

36. Robberecht C VT, Zamani Esteki M, Nowakowska BA, Vermeesch JR. Non-allelic homologous recombination between retrotransposable elements is a driver of de novo unbalanced translocations. Genome Res. 2013;23:411–18.

37. Cai X, Xu SS. Meiosis-driven genome variation in plants. Curr Genomics. 2007;8:151–61.

38. Storici F, Snipe JR, Chan GK, Gordenin DA, Resnick MA. Conservative repair of a chromosomal double-strand break by single-strand DNA through two steps of annealing. Mol Cell Biol. 2006;6:7645–57.

39. Yang Y, Sterling J, Storici F, Resnick MA, Gordenin DA. Hypermutability of damaged single-Strand DNA formed at double-strand breaks and uncapped telomeres in yeast *Saccharomyces cerevisiae*. Plos Genet. 2008;4:e1000264.

40. Fishman-Lobell JRN, Haber JE. Two alternative pathways of double-strand break repair that are kinetically separable and independently modulated. Mol Cell Biol. 1992;12:1292–1303.

41. Shevelev IV, Hubscher U. The 3 ‘-5’ exonucleases. Nat Rev Mol Cell Biol. 2002;3:364–75.

42. Pfeiffer P, Goedecke W, Obe G. Mechanisms of DNA double-strand break repair and their potential to induce chromosomal aberrations. Mutagenesis. 2000;15:289–302.

43. Leitch AR, Leitch IJ. Genomic plasticity and the diversity of polyploid plants. Science. 2008;320:481–3.

44. Dyck PL, Kerber ER. Inheritance in hexaploid wheat of adult-plant leaf rust resistance derived from *Aegilops squarrosa*. Can J Genet Cytol. 1970;2:175–80.

45. Wang J, Luo MC, Chen ZX, You FM, Wei YM, Zheng YL, Dvorak J. *Aegilops tauschii* single nucleotide polymorphisms shed light on the origins of wheat D-genome genetic diversity and pinpoint the geographic origin of hexaploid wheat. New Phytol. 2013;198:925–37.

46. Arora S, Singh N, Kaur S, Bains NS, Uauy C, Poland J, Chhuneja P. Genome-wide association study of grain architecture in wild wheat *Aegilops tauschii*. Front Plant Sci. 2017;8:886.

47. Luo MC, Gu YQ, Puiu D, Wang H, Twardziok SO, Deal KR, Huo N, Zhu T, Wang L, Wang Y, et al. Genome sequence of the progenitor of the wheat D genome *Aegilops tauschii*. Nature. 2017;551:498–502.

48. Genetic Resources Information System for Wheat and Triticale. http://www.wheatpedigree.net/sort/show/118822. Accessed 19 Dec 2017.

49. Isidore E, Scherrer B, Chalhoub B, Feuillet C, Keller B. Ancient haplotypes resulting from extensive molecular rearrangements in the wheat A genome have been maintained in species of three different ploidy levels. Genome Res. 2005;15:526–36.

50. Saxena RK, Edwards D, Varshney RK. Structural variations in plant genomes. Brief Funct Genomics. 2014;13:296–307.

51. Mago R, Tabe L, Vautrin S, Simkova H, Kubalakova M, Upadhyaya N, Berges H, Kong X, Breen J, Dolezel J, et al. Major haplotype divergence including multiple germin-like protein genes, at the wheat *Sr2* adult plant stem rust resistance locus. BMC Plant Biol. 2014;14:379.

52. Pearce S, Zhu J, Boldizsar A, Vagujfalvi A, Burke A, Garland-Campbell K, Galiba G, Dubcovsky J. Large deletions in the CBF gene cluster at the *Fr-B2* locus are associated with reduced frost tolerance in wheat. Theor Appl Genet. 2013;126:2683-97.

53. Chia JM, Song C, Bradbury PJ, Costich D, de Leon N, Doebley J, Elshire RJ, Gaut B, Geller L, Glaubitz JC, et al. Maize HapMap2 identifies extant variation from a genome in flux. Nat Genet. 2012;44:803–7.

54. The 3,000 rice genomes project. The 3,000 rice genomes project. Gigascience. 2014;3:7.

55. International Rice Genome Sequencing Project. The map-based sequence of the rice genome. Nature. 2005;436:793–800.

56. Paterson AH, Bowers JE, Bruggmann R, Dubchak I, Grimwood J, Gundlach H, Haberer G, Hellsten U, Mitros T, Poliakov A, et al. The *Sorghum bicolor genome* and the diversification of grasses. Nature. 2009;457:551–6.

57. Schnable PS, Ware D, Fulton RS, Stein JC, Wei F, Pasternak S, Liang C, Zhang J, Fulton L, Graves TA, et al. The B73 maize genome: complexity, diversity, and dynamics. Science. 2009;326:1112–5.

58. Vaughn JN, Bennetzen JL. Natural insertions in rice commonly form tandem duplications indicative of patch-mediated double-strand break induction and repair. Proc Natl Acad Sci USA. 2014;111:6684–9.

59. Buchmann JP, Matsumoto T, Stein N, Keller B, Wicker T. Inter-species sequence comparison of Brachypodium reveals how transposon activity corrodes genome colinearity. Plant J. 2012;71:550–63.

60. Woodhouse MR, Schnable JC, Pedersen BS, Lyons E, Lisch D, Subramaniam S, Freeling M. Following tetraploidy in maize, a short deletion mechanism removed genes preferentially from one of the two homeologs. PLoS Biol. 2010;8:e1000409.

61. Sudupak MA, Bennetzen JL, Hulbert SH. Unequal exchange and meiotic instability of disease-resistance genes in the *RP1* region of maize. Genetics. 1993;133:119–25.

62. Ramakrishna W, Emberton J, Ogden M, SanMiguel P, Bennetzen JL. Structural analysis of the maize *RP1* complex reveals numerous sites and unexpected mechanisms of local rearrangement. Plant Cell. 2002;14:3213–23.

63. Sandhu D, Gao HY, Cianzio S, Bhattacharyya MK. Deletion of a disease resistance nucleotide-binding-site leucine-rich-repeat-like sequence is associated with the loss of the Phytophthora resistance gene *Rps4* in soybean. Genetics. 2004;168:2157–67.

64. Hiebert CW, Thomas JB, Somers DJ, McCallum BD, Fox SL. Microsatellite mapping of adult-plant leaf rust resistance gene *Lr22a* in wheat. Theor Appl Genet. 2007;115:877–84.

65. Tanksley SD, McCouch SR. Seed banks and molecular maps: unlocking genetic potential from the wild. Science. 1997;277:1063–6.

66. Mcfadden ES, Sears ER. The artificial synthesis of Triticum-Spelta. Records Genet Soc Amer. 1944;13:26–7.

67. Dreisigacker S, Kishii M, Lage J, Warburton M. Use of synthetic hexaploid wheat to increase diversity for CIMMYT bread wheat improvement. Aus J Agric Res. 2008;59:413–20.

68. Molnár-Láng M, Ceoloni C, Dolezel J. Alien introgression in wheat: Cytogenetics, molecular biology, and genomics. Springer International Publishing; 2015.

69. Greenwood TA, Rana BK, Schork NJ. Human haplotype block sizes are negatively correlated with recombination rates. Genome Res. 2004;14:1358–61.

70. Denton JF, Lugo-Martinez J, Tucker AE, Schrider DR, Warren WC, Hahn MW. Extensive error in the number of genes inferred from draft genome assemblies. PLoS Comput Biol. 2014;10:e1003998.

71. Zapata L, Ding J, Willing EM, Hartwig B, Bezdan D, Jiao WB, Patel V, Velikkakam James G, Koornneef M, Ossowski S, Schneeberger K. Chromosome-level assembly of *Arabidopsis thaliana Ler* reveals the extent of translocation and inversion polymorphisms. Proc Natl Acad Sci USA. 2016;113:e4052–60.

72. Gremme G, Brendel V, Sparks ME, Kurtz S. Engineering a software tool for gene structure prediction in higher organisms. Inform Software Tech. 2005;47:965–78.

73. NLR Annotator: https://github.com/steuernb/NLR-Annotator. Accessed 4 Jan 2018.

74. MicrobesOnline: http://www.microbesonline.org/fasttree/. Accessed 21 feb 2017.

75. Price MN, Dehal PS, Arkin AP. FastTree 2—approximately maximum-likelihood trees for large alignments. PLoS One. 2010;5:e9490.

76. Sonnhammer EL, Durbin R. A dot-matrix program with dynamic threshold control suited for genomic DNA and protein sequence analysis. Gene. 1995;167:GC1–10.

77. Altschul SF, Madden TL, Schaffer AA, Zhang J, Zhang Z, Miller W, Lipman DJ. Gapped BLAST and PSI-BLAST: a new generation of protein database search programs. Nucleic Acids Res. 1997;25:3389–402.

78. The International Brachypodium Initiative. Genome sequencing and analysis of the model grass *Brachypodium distachyon*. Nature. 2010;463:763–8.

